## Supplementary material for "G2PDeep-v2: a web-based deep-learning framework for phenotype prediction and biomarker discovery using multi-omics data": supplementary Fig. S1.docx

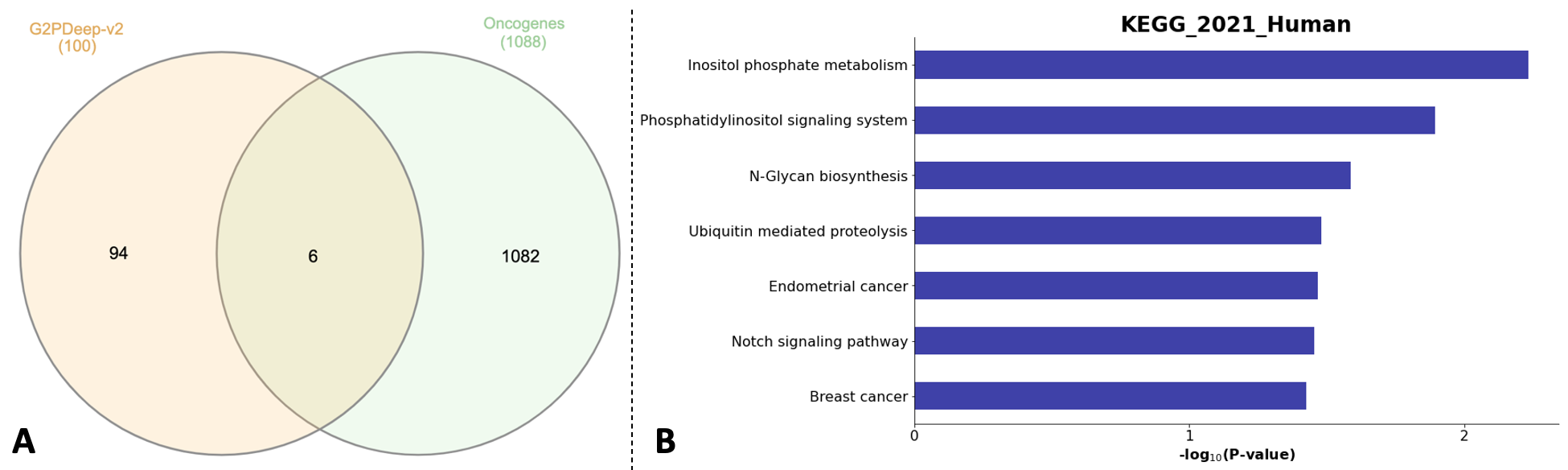

**Supplementary Fig. 1**. Plots for significant biomarkers. (A) Venn diagram of 100 most significant biomarkers and oncogenes. (B) Enriched seven pathways associated with breast cancer.
